# Data-driven evaluation of suitable immunogens for improved antibody selection

**DOI:** 10.1101/2024.10.07.617016

**Authors:** Katharina Waury, Hlin Kvartsberg, Henrik Zetterberg, Kaj Blennow, Charlotte E. Teunissen, Sanne Abeln

## Abstract

Antibodies are indispensable in laboratory and clinical applications due to their high specificity and affinity for protein antigens. However, selecting the right protein fragments as immunogens for antibody production remains challenging. Leveraging the Human Protein Atlas, this study systematically evaluates immunogen properties aiming to identify key factors that influence their suitability. Antibodies were classified as successful or unsuccessful based on standardized validation experiments, and the structural and functional properties of their immunogens were analyzed. Results indicated that longer immunogens often resulted in more successful but less specific antibodies. Shorter immunogens (50 residues or fewer) with disordered or unfolded regions at the N- or C-terminus and long coil stretches, were more likely to generate successful antibodies. Conversely, immunogens with high beta sheet content, multiple secondary structure elements, transmembrane regions, or disulfide bridges were associated with poorer antibody performance. Post-translational modification sites within immunogens appeared to mark beneficial regions for antibody generation. To support antibody selection, a novel R package, immunogenViewer, was developed, enabling researchers to easily apply these insights when immunogen sequences are disclosed. By providing a deeper understanding of immunogen suitability, this study promotes the development of more effective antibodies, ultimately addressing issues of reproducibility and reliability in antibody-based research. The findings are highly relevant to the research community, as end-users often lack control over the immunogen selection process in antibody production.

## Introduction

Antibodies are among the most essential tools for sensitive and reliable protein detection in both laboratory and clinical settings^1,2^. Their specificity and affinity make them invaluable in many molecular biology-based technologies for the detection and quantification of proteins, such as western blots (WBs) and immunohistochemistry (IHC)^3^. These technologies are fundamental to both basic and applied scientific investigations, providing critical insights into biological processes and disease mechanisms. In the clinic, antibodies play a pivotal role in immunoassays to facilitate robust biomarker detection for diagnosis, prognosis, and patient stratification^4,5^.

Antibodies are typically produced through the immunization of animals with an antigen, a process that generates a robust immune response against the target protein^6^. Polyclonal antibodies can subsequently be purified from the serum of the immunized animal. While other approaches are available, e.g., hybridoma cell cultures, animal immunization offers many advantages. This technology is very well established and widespread^7^. Further, antibodies derived from animal immunization often outperform those produced by in vitro display technologies regarding affinity and specificity^6,7^.

Before the immunization of the animal, an essential step is the selection of the type of immunogen. The immunogen is the molecule injected into the animal to induce an immune response that leads to the production of the desired antibodies^8^. When the aim is to produce antibodies recognizing a specific protein, several options are available regarding the immunogen. Other than the full-length protein, using only one domain, a shorter fragment of the protein, or even a peptide can be considered^9^. There are multiple reasons for choosing a protein fragment or peptide instead of a full-length, natively folded protein. It may be challenging or impossible to produce and purify full proteins due to their size, complexity, or instability^10,11^. In such cases, peptide production is more straightforward and can be easily outsourced^10^. Additionally, the use of a peptide enables the generation of isoform- or proteoform-specific antibodies to distinguish between closely related protein variants^12^.

Using protein fragments or peptides for antibody generation also allows researchers to more precisely locate the epitope, which is the antibody recognition site on the protein. By selecting specific peptide sequences, the resulting antibodies already have fairly well-characterized epitopes and known specificity^12^. This can be very beneficial when developing a sandwich immunoassay. As in this application two different antibodies bind the desired protein simultaneously, it is vital that these two antibodies do not have the same or overlapping recognition site unless a homomeric protein is to be detected^13^.

While offering multiple advantages, the selection of the most suitable peptide or protein fragment is a highly complex and critical task^10^. Especially if the antibody is raised against a peptide but intended to detect the natively folded form of the protein, identifying the most appropriate immunogen sequence is vital to produce working antibodies^11^. Brown *et al.* compared the performance of antibodies raised against peptides and full-length proteins^9^. In their study, anti-peptide antibodies performed worse, especially in applications which required native protein detection such as enzyme-linked immunosorbent assays (ELISAs). Several potential pitfalls can compromise the efficacy of antibodies resulting from immunization with peptides or protein fragments:

- Surface accessibility: The selected peptide may not be located at the surface in the folded protein, hindering accessibility and thus antibody binding^14^.
- Structural conformation: The peptide in isolation might adopt a different structural conformation than when it is incorporated within the full-length protein sequence. Thus, the immunogen against which the antibody was raised would not be recognizable within the protein and negatively affect antibody function^10,11^.
- Specificity: The peptide may not be unique to the target protein, leading to cross-reactivity with other antigens that share the same sequence^10^.

While guidelines^10,11,14^ exist to aid in the selection of suitable peptides or protein fragments, these suggestions are often not quantified or supported by extensive data. This lack of robust, data-driven guidelines can lead to inconsistencies in antibody performance and reliability and incomplete leverage of knowledge. Recommendations include the selection of protein sequences with a high loop or turn content, high surface accessibility, and hydrophilicity^10,11,14^. It has also been suggested to choose immunogens that are located within the N- or C-terminus of the protein sequence as these regions are often exposed and flexible^10,11,14^. Note that hereinafter we solely focus on the suitability of immunogens in regard to facilitating antibody recognition. The aspect of immunogenicity, i.e., the ability of a molecule to elicit an immune response, is not considered explicitly.

To address the challenges of the most suitable immunogen choice, our study aims to systematically analyze antibodies with known immunogens to understand the most optimal characteristics of these protein fragments. For this purpose, we utilized the Human Protein Atlas (HPA)^15^, a unique and highly valuable data resource providing immunogen information for their in-house produced polyclonal antibodies. Selection of the immunogens reported in the HPA already incorporates important factors including limited sequence similarity to the human proteome, and the absence of transmembrane regions and signal peptides^16,17^. Further, antibodies are tested and validated in a highly consistent manner. Results for three types of antibody-based technologies are contained in the atlas: 1) WBs allow protein identification based on molecular weight, 2) IHC uses immunostaining to localize proteins within tissue samples, 3) Immunocytochemistry (ICC) identifies subcellular protein localization within cells^3^. Importantly, negative results are also published in the atlas^15,18^. This allows us to define “successful” antibodies if they work in the majority of these three technologies, i.e., WB, IHC and ICC. Vice versa, “unsuccessful” antibodies are identified based on the majority of the technologies failing.

Identifying relevant properties for immunogen evaluation with this data-driven approach is extremely relevant to the research community considering that many molecular biology-based technologies are developed using commercial antibodies where the end-user cannot influence the choice of the immunogen sequence^9^. Choosing the most appropriate and reliable antibody for the intended application is an important step in overcoming the current reproducibility and reliability issues of research using antibody technologies due to hindered or unspecific antibody binding^19^. Finally, to allow researchers to quickly and easily use the insights gained in this work in their antibody or immunogen selection process, we developed the novel open access R package *immunogenViewer* to support immunogen evaluation and antibody selection.

## Results

To allow analysis of the properties of protein fragments suitable to be used as immunogens, the HPA was mined for relevant information regarding in-house antibodies, most importantly their performance success across three different antibody-based applications and their respective immunogen. This data allowed the annotation of antibodies as unsuccessful or successful and the comparison of their respective immunogen sequences. Annotations were generated, including AlphaFold2-based structural features and curated annotations from the UniProt database, and compared to identify which properties are most important when selecting immunogen sequences.

### The Human Protein Atlas as a suitable data source to evaluate immunogen suitability

Data mining of the atlas led to a dataset of 23.616 antibodies. The HPA antibodies cover 80.79% of the human proteome (Figure 1A). Of the proteins of the human proteome according to UniProt, 9210 proteins (45.07%) had one antibody targeting it, 7300 proteins (35.72%) had two or more antibodies targeting it.

**Figure 1.**
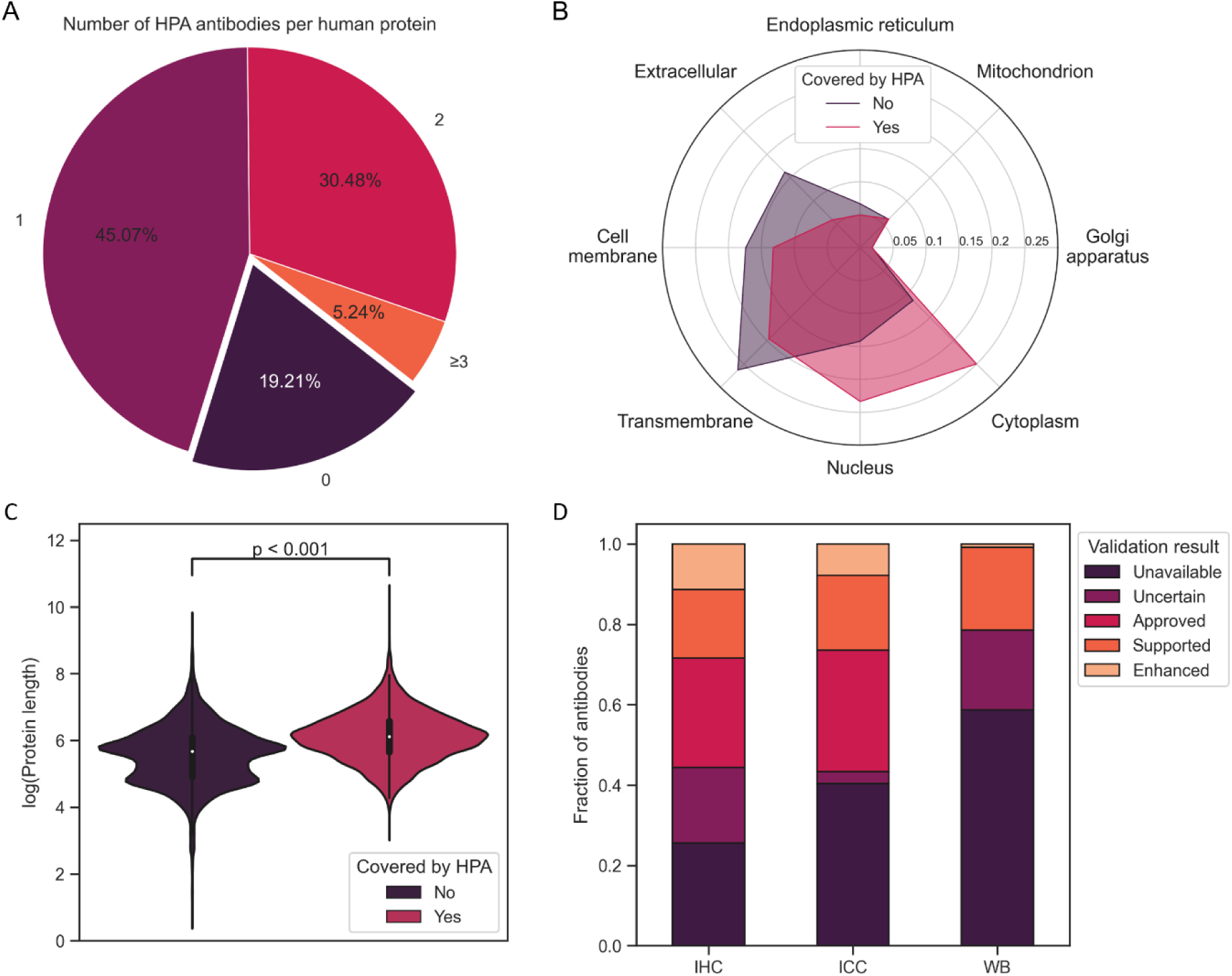
Coverage of the Human Protein Atlas. Validated HPA antibodies cover more than 80% of the human proteome. More than one third of the proteins are associated with multiple antibodies (A). The subset of the proteome covered by the HPA contains more proteins predicted to be located within the cytoplasm and nucleus. In comparison, proteins for which no HPA antibody is available are more likely to be at the cell membrane or in the extracellular space, and contain transmembrane regions (B). The proteome not covered by the HPA is enriched in smaller proteins with shorter sequence length (C). The three included antibody-based methods (IHC, ICC and WB) show different coverage and success rates. The most antibodies have been established in IHC, ICC is the most successful method. For less than half of the HPA antibodies WB results are available (D). HPA - Human Protein Atlas; ICC – Immunocytochemistry; IHC – Immunohistochemistry; WB - western blot

The part of the proteome for which no validated antibodies are available was enriched in transmembrane proteins and proteins associated with the cell membrane or the extracellular space (Figure 1B). Visualization of the protein sequence lengths of covered and not covered proteins indicated a subset of smaller proteins for which no HPA antibodies are available (Figure 1C). A possible reason is that the immunogen selection algorithm applied for HPA antibody generation^16^ was not able to find a protein fragment with low enough sequence identity to the human proteome in these short proteins.

According to the HPA, antibodies are categorized as “uncertain” for a tested technology (WB, IHC or ICC), if validation experiments show undesired or differing results compared to mRNA analysis or literature. Antibodies working as intended are labelled “approved”, “supported”, or “enhanced” depending on the level of evidence and validation^20^. Comparison of the antibody validation results showed that coverage is not equal across the three tested technologies. IHC had the highest coverage across all HPA antibodies (74.43%), while ICC had the highest success rate across all HPA antibodies for which this technology had been tested (95.06%). For more than half of the HPA antibodies WB results were not available. Note that while more WB results are deposited in the atlas, only WB results with an associated image could be accessed in a high-throughput manner. Based on these validation results in WB, IHC and ICC, HPA antibodies and their respective immunogen were classified as “successful” or “unsuccessful” (see Methods).

### Longer immunogens increase antibody success rate but are less specific

The included successful and unsuccessful HPA antibodies were raised against immunogens ranging in length from 19 to 202 residues. We compared the effect of increasing immunogen length both on antibody success rate and WB specificity. Longer immunogens generated successful antibodies more often than shorter immunogens. While the 707 immunogens below a length of 30 residues showed a success rate of 78.08%, this increased steadily with a growing immunogen length to a success rate of 93.68% in antibodies that were raised against immunogens of 110 residues or more (Figure 2A). On the other hand, including information on the antibody specificity highlighted that off-target binding in WBs becomes more likely in longer immunogens (Figure 2B). Considering that HPA antibodies are polyclonal, i.e., the antibodies can have differing binding regions against the utilized immunogen, these observations were to be expected. A longer immunogen provides more potential binding regions to raise working antibodies against^21^ but might also contain regions that are similar to other protein sequences of the human proteome leading to off-target binding of the respective antibodies.

**Figure 2.**
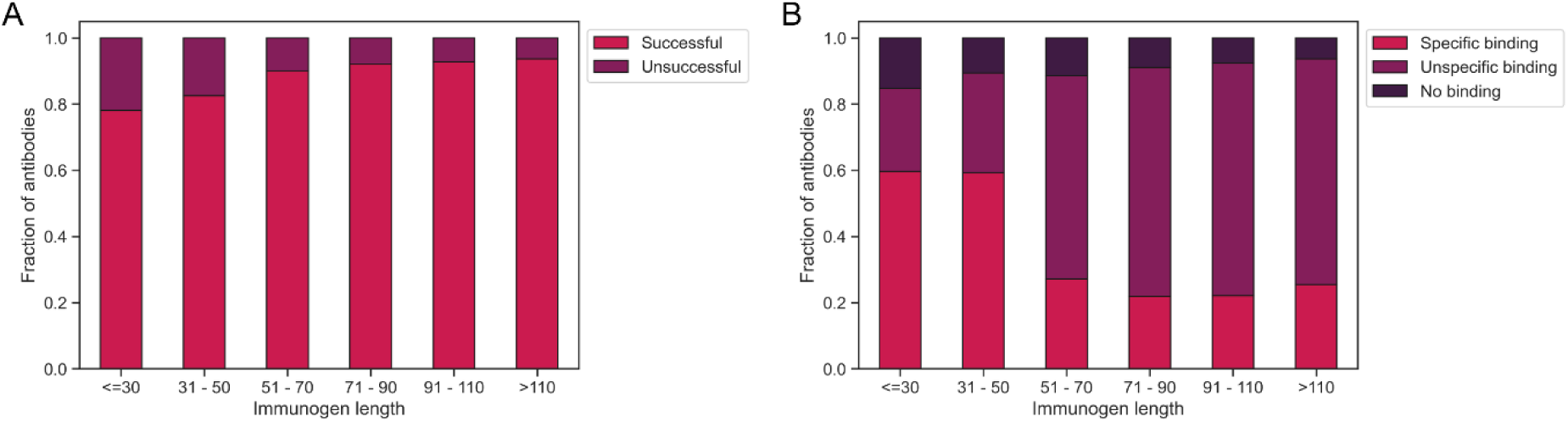
Antibody success across immunogen length. The success rate of HPA antibodies (defined by the results of validation experiments) is continuously higher with increasing length of the immunogen used for antibody generation (A). While the fraction of unsuccessful antibodies, decreases in WB experiments, unspecific binding is more commonly observed in antibodies raised against longer immunogens (B). HPA - Human Protein Atlas; WB - western blot

In the following, we focused only on immunogens comprised of 50 residues or less. This filter criterion still provides us with a dataset of 815 successful and 188 unsuccessful antibodies and their respective immunogens, while the antibody-binding regions were better defined because of the limited immunogen size allowing a more concrete analysis of immunogen suitability. This is especially relevant considering that HPA antibodies are polyclonal, and an increasing immunogen length leads to a higher number of possible epitopes and thus more diversity in the antibodies being generated^22^.

### Disordered terminus regions are confirmed as advantageous immunogen regions

Consistently mentioned properties to consider during immunogen selection are flexibility and surface accessibility^10,11,14^. Protein disorder provides a good proxy for flexible loop regions^23^ and intrinsically disordered regions are generally known to have high solvent accessibility^24^. We thus wished to investigate if disorder is a strong indication of immunogen suitability. Furthermore, as N-and C-terminal regions have been recommended specifically as suitable immunogen regions^10^, we further divided our immunogens based on their position within the full protein sequence. An immunogen’s position was designated as “terminus” if located at least partly within the first or last 25 residues of the protein sequence, otherwise the position was considered “center”. These definitions separated the immunogens relatively equally into two (Table 1). Interestingly, within the HPA data we did not observe a large difference in success rate between terminus and center immunogens (Table 1).

**Table 1.** Dataset compositions in regard to immunogen location within the protein sequence.

| Position | Number | Success rate |
| --- | --- | --- |
| Terminus | 459 | 81.92% |
| Center | 538 | 80.48% |

We used predicted disorder scores of each immunogen residue to define 398 low, 491 medium, and 114 high disorder immunogens (see Methods). Comparison of success rates in these disorder groups is shown in Figure 3A. Two trends became apparent: 1) disorder increased success rates for immunogens regardless of their positions; 2) the increased disorder content had a stronger effect in terminus immunogens. In this group, we saw an increase in the antibody success rate from low to high disorder immunogens of 35.98%, compared to an increase of only 10.65% when immunogens are located within the center.

**Figure 3.**
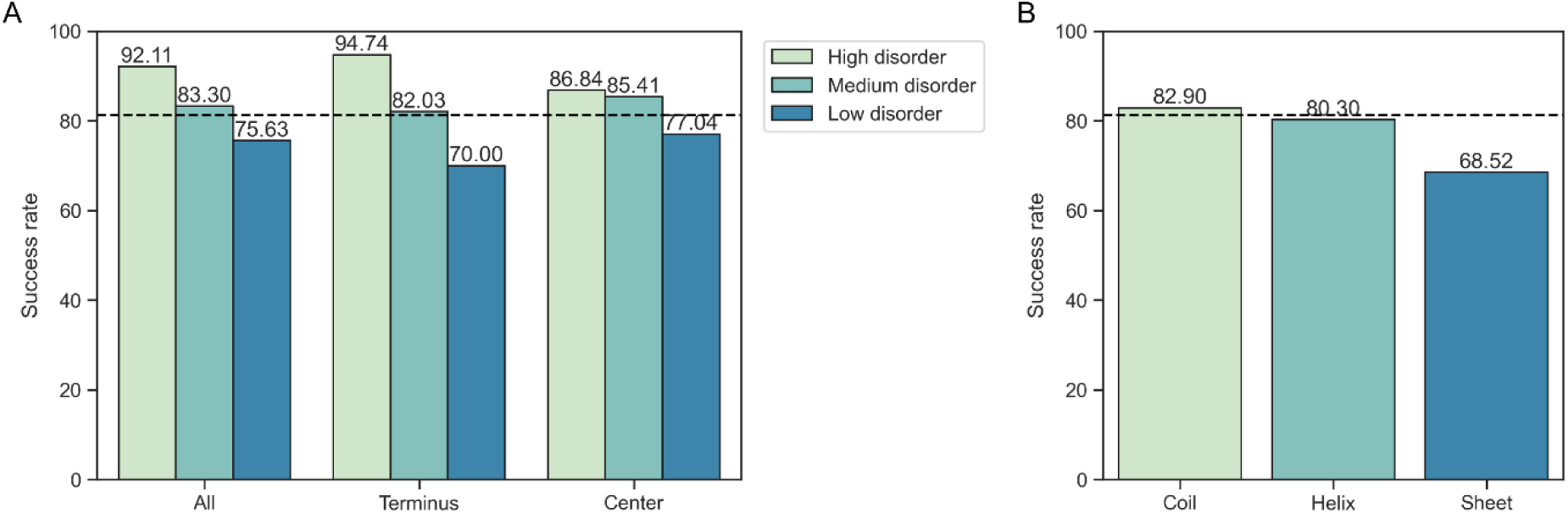
Antibody success rate in regard to disorder and secondary structure. Highly disordered immunogens are more likely to raise working antibodies. The positive effect of disorder on antibody success is more pronounced when the immunogen is located within the N- or C-terminus of the protein sequence. Highly disordered immunogen within the terminus have the highest success rate. The difference in success rates is significant across all three groups (p_All_ < 0.001, p_Terminus_ < 0.001, p_Center_ = 0.044) (A). The most prevalent secondary structure of immunogen residues, i.e., the majority secondary structure, influences antibody success rates significantly (p = 0.03). Coil residues are advantageous, sheet residues are the most detrimental (B). The dashed black line indicates the success rate across the full antibody set.

In conclusion, high disorder terminal immunogens showed the highest success rates suggesting that these regions are suitable immunogens.

### High sheet content in immunogens is detrimental

Regions that do not form alpha helices or beta sheets, e.g., connecting loop or turn regions, have been suggested to be superior immunogen choices^11^. We refer to all regions that do not adopt a helix or sheet conformation as coils hereinafter. We wished to compare the impact of secondary structure on immunogen suitability as differences exist between them regarding important properties such as accessibility and stability^25,26^. Residues in every immunogen were assigned as coil, helix or sheet based on all proteins’ predicted AlphaFold2 structures^27^. We then identified the majority secondary structure, i.e., the type that is most prevalent, for each immunogen and compared their success rates. Immunogens dominated by coils led most often to successful antibodies (82.82%), closely followed by immunogens prevalent in helix residues (80.12%). There was a clear drop in the success rate (68.52%) when the majority of immunogen residues take a sheet conformation (Figure 3B).

Most immunogens fell into the coil majority class, while only a small number of immunogens were assigned to the sheet majority class. Comparison of the average sheet content of the full protein sequences in our dataset with the sheet content of the immunogen sequences showed that immunogens in the HPA seem to be biased towards protein fragments devoid of sheets (Figure S1A). Furthermore, in the immunogens with beta sheets being the majority secondary structure many residues still adopted a coil or helix conformation. Immunogens in which coils or helices were most prevalent, often no other conformation was found (Figure S1B). These results suggest that a high content of beta sheets within immunogens is a potential obstacle for generating functioning antibodies. This issue would likely be more detrimental if no other secondary structure stretch is present that can serve as a recognition site for antibodies.

### Other structural differences between successful and unsuccessful immunogens

Another investigated immunogen feature is the number of distinct secondary structure elements within the immunogen sequence. Interestingly, a difference is only observed between unsuccessful and successful immunogens regarding the number of changes between secondary structures in center-located immunogens (Figure 4A). This might indicate that generation of working antibodies is more likely to succeed when longer stretches of one secondary structure are available instead of alternating shorter secondary structure elements.

**Figure 4.**
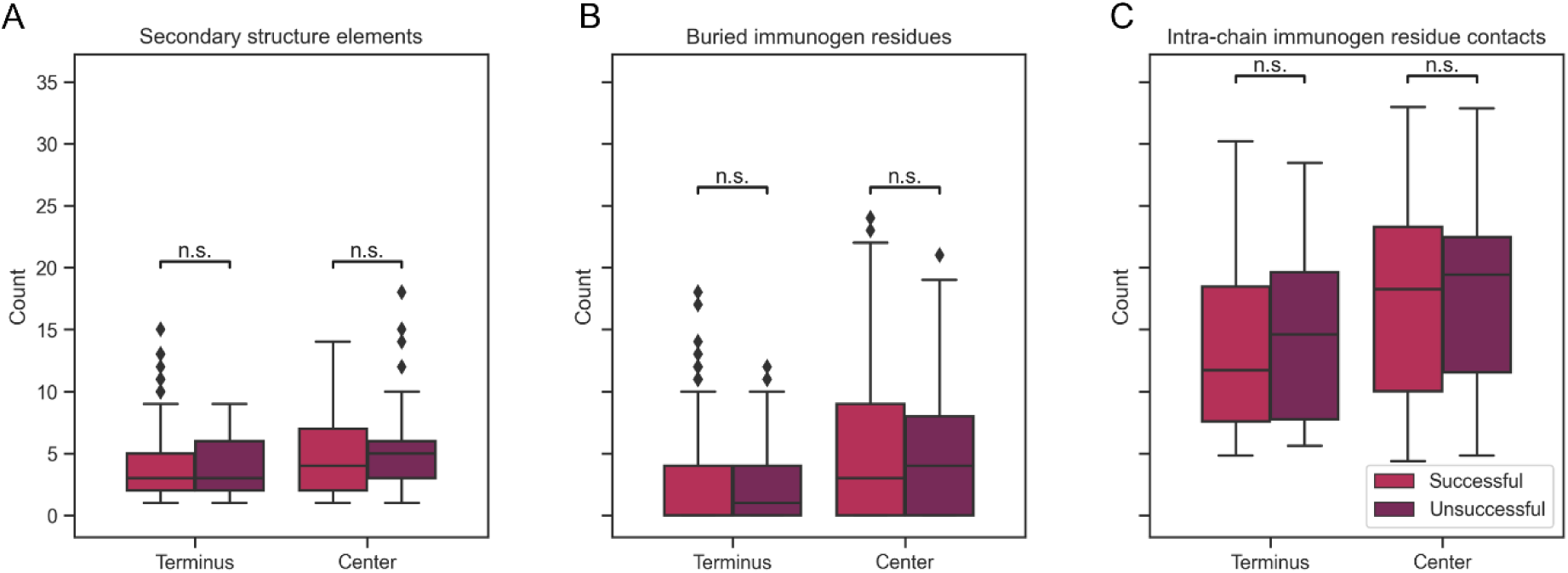
Structural differences between successful and unsuccessful immunogens. Changes between secondary structures within the immunogen sequence are more common in unsuccessful immunogens when located towards the center of the protein sequence (A). Buried residues have a very low relative surface accessibility as these are buried within the protein core and are not available as binding regions. Center-located immunogens contain more buried residues. The number of buried residues is higher in unsuccessful immunogen sequences (B). The number of intra-chain contacts per residue can provide an estimate of the degree of folding within a protein. A higher number of contacts per residues is associated with unsuccessful immunogens (C). Because of limited effect and dataset size none of these differences are statistically significant (p > 0.05).

Two further structural properties gave an indication about the accessibility of the immunogen and the fold of the protein. The number of buried residues measures what part of the immunogen is truly inaccessible to antibody binding within the natively folded protein structure (Figure 4B). The average number of intra-chain contacts provided information about how strong the fold of the protein is within the area of the immunogen. A higher number of intra-chain contacts suggests that many residues are within proximity of the immunogen residues, as is the case in globular folded proteins and core residues^28^ (Figure 4C). Both of these features generally had a higher number in our dataset within the protein center, as terminus regions are often flexible and not folded. Across both terminus and center immunogens we observed that a higher degree of protein folding, i.e., a lower degree of solvent accessibility, was associated with unsuccessful antibodies.

Taken together, indications of the immunogen being located within a tightly folded region of the protein sequence should be avoided during immunogen selection (Figure 4). Note however that the effect sizes are not large suggesting that no single structural feature could provide enough information to judge an immunogen as suitable or unsuitable.

### Transmembrane regions, disulfide bridges, PTMs

In addition to an analysis of structural features, we were interested to identify if functional annotations had an impact on immunogen suitability. We investigated if the presence of transmembrane regions, disulfide bridges and post-translational modification (PTM) sites in immunogens affected the generation of successful antibodies.

Through a protein’s incorporation into the cell membrane part of it becomes inaccessible and thus not suitable for binding of an antibody^16^. Thus as is expected, the presence of a transmembrane region within the protein to be detected led to a lower percentage of successful antibodies (Figure 5A). This observation is in line with our earlier results that proteins not covered by any HPA antibodies were enriched in transmembrane regions (Figure 1B), suggesting that for these proteins no validated antibody could be generated. The presence of transmembrane regions was most unfavourable if immunogen residues overlapped with the transmembrane region, compared to immunogens that were only in the vicinity of the membrane-bound part of the protein.

**Figure 5.**
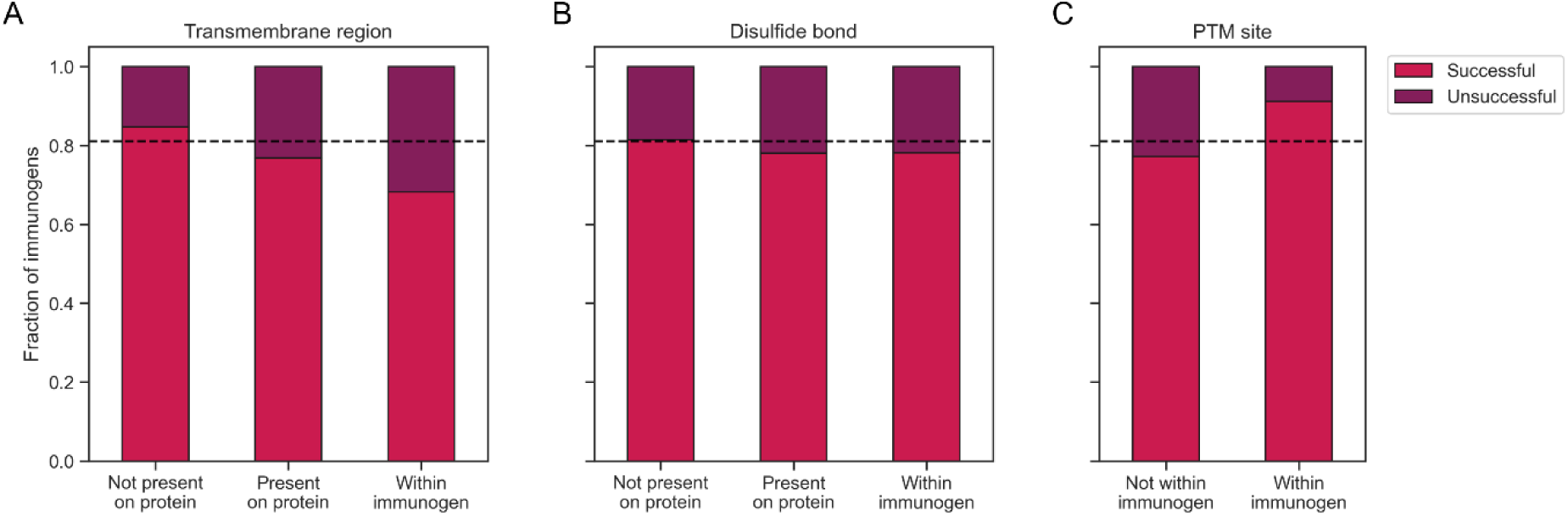
Functional annotations of immunogens. Presence of a transmembrane region within the protein antigen lowers the success rate of antibodies. The success rate is significantly lower when the immunogen used to raise antibodies overlaps with the transmembrane region (p < 0.001) (A). The presence of disulfide bridges lowers the success rate of antibodies, but the effect is less pronounced compared to transmembrane regions (p = 0.687) (B). The presence of PTM sites within the immunogen is more common in successfully working antibodies (p < 0.001). Many PTM sites have to be accessible and are thus also available for binding. The PTM annotation does not guarantee that this site is modified when testing the HPA antibodies (C). The dashed black line indicates the success rate across the full antibody set. HPA - Human Protein Atlas; PTM - post-translational modification

Disulfide bridges might also indicate unfavourable immunogen sequences as these tend to be buried^29^ and thus the immunogen itself might be in an inaccessible position within the full protein. While we also saw a lower success rate for immunogens in which a disulfide bridge is present, the difference was less pronounced in this dataset compared to the presence of transmembrane regions (Figure 5B). Note that disulfide bridge annotations are more scarce than transmembrane regions (103 proteins with a disulfide bond vs. 318 transmembrane proteins).

Finally, we compared the effect of PTM sites within the immunogens on antibody success. Here, annotations of modified sites were more common within successful immunogens (Figure 5C). While PTMs have been suggested to be a potential obstacle to antibody binding if the immunogen used to raise antibodies did not contain the modification as well^13^, here we saw a beneficial impact of modified sites on antibody success. It is important to consider, however, that available PTM annotations only provide information on residues that are known to have been modified. This is not direct evidence that any PTMs were actually present in the experiments done within the HPA project. As most PTM sites need to be surface accessible for a modification to be added to the relevant residue by an enzyme and for the modification to perform its function^30^, PTM sites could thus be interpreted as another proxy for residue accessibility.

### Fast and effortless immunogen evaluation with immunogenViewer

The comprehensive analysis of successful and unsuccessful immunogens led to a list of properties that should be considered when selecting novel or judge known immunogens. To facilitate an easy manner in which to evaluate the suitability of protein fragments or peptides as immunogens, we developed the R package *immunogenViewer*. This tool allows the visualization of immunogens of interest along the protein sequence and highlights relevant properties to consider including disorder, secondary structure, surface accessibility and PTMs. Importantly, this tool has a broad applicability as it can be used for any human protein. The package comes with a detailed manual and tutorial allowing researcher to use the package for their own needs. The package *immunogenViewer* can be accessed freely and installed easily via Bioconductor. All necessary information can be found at https://bioconductor.org/packages/devel/bioc/html/immunogenViewer.html. As an example, the predicted structure of the human protein Histone deacetylase complex subunit SAP30L (Figure 6A) and the visualization of its protein properties using *immunogenViewer* (Figure 6B) are shown with the immunogens of two HPA antibodies highlighted both along the structure and the sequence. Both antibodies were classified as successful based on the validation results of the HPA. The respective immunogens show a high fraction of disordered residues within the N- and C-terminus which we identified as a suitable property in this work.

**Figure 6.**
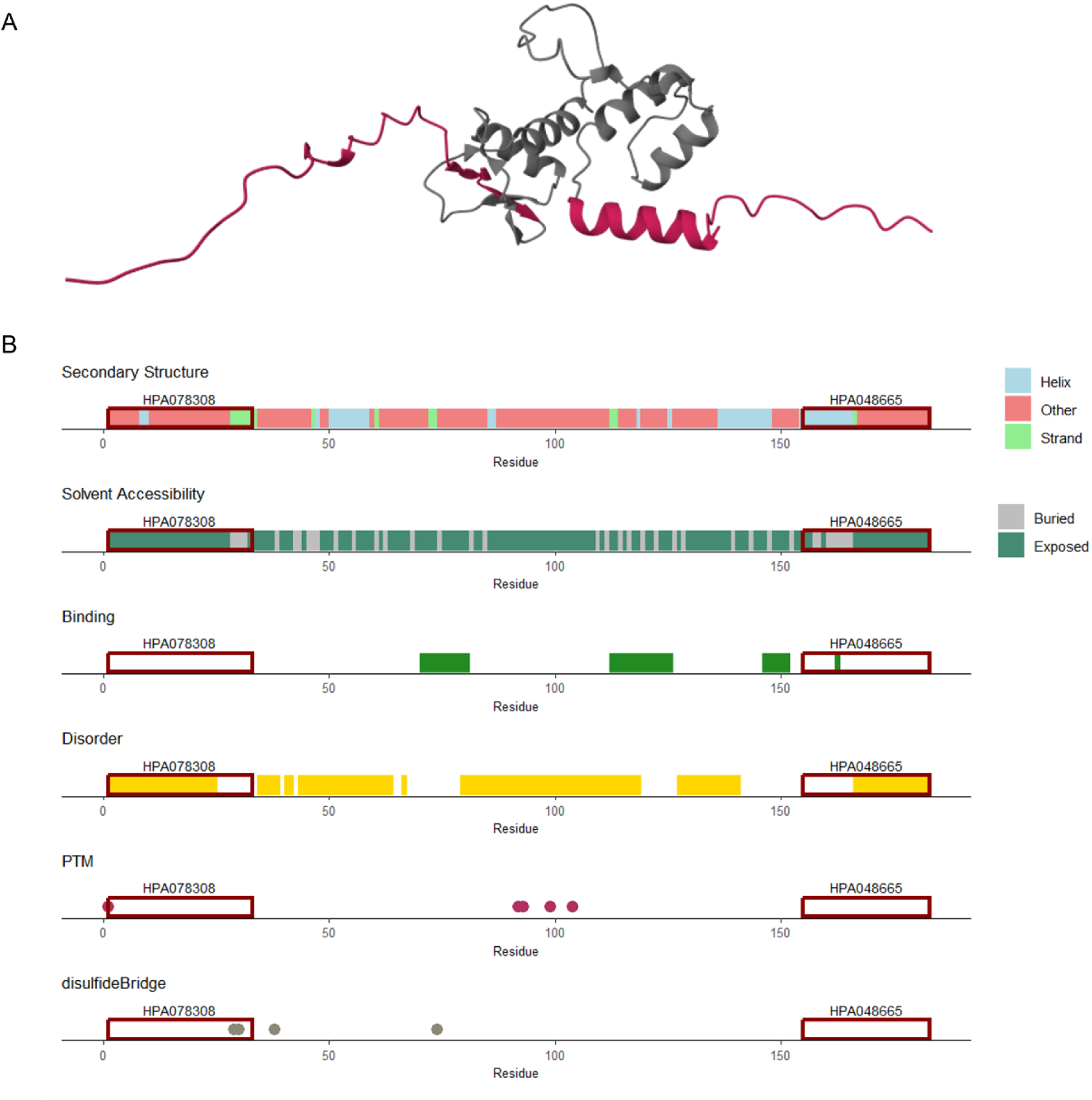
Protein visualization and immunogen evaluation with *immunogenViewer*. Protein structure of Histone deacetylase complex subunit SAP30L (UniProt ID: Q9HAJ7) according to the AlphaFold Protein Structure Database (A). Output of *immunogenViewer* visualizing several properties along the protein’s sequence. Two available HPA antibodies are highlighted which can be evaluated using the package (B). HPA - Human Protein Atlas; PTM - post-translational modification

## Discussion

Knowledge of suitability of immunogens and their respective antibodies is strongly needed as available guidelines on immunogen selection are scarce and not data-driven. Our analysis aimed to address this gap, leading to the confirmation of existing recommendations such as choosing terminus and coil regions. We expanded the knowledge of optimal immunogen and antibody selection by comparing additional structural and functional features. Importantly, the relevant properties identified here can be evaluated easily and rapidly using our developed R package *immunogenViewer*.

Our study shows that properties increasing the suitability of immunogens to generate functioning antibodies include a high disorder content, especially within the terminus regions, and uninterrupted coil or helix stretches that are solvent accessible. Detrimental properties include a higher number of distinct secondary structure elements, as well as a high number of intra-chain residue contacts and buried residues. High content of beta sheets is also associated with lower success rates. In addition to the generally lower solvent accessibility of beta sheets^26^, another reason for this observation could be the non-local hydrogen bonds that are formed between beta sheets^31^. We hypothesize that protein regions with a sheet conformation within the full-length protein may be problematic as this conformation might differ within the immunogen in isolation if the necessary hydrogen-binding partners are not available, in addition to immunogen accessibility.

The presence of transmembrane regions and disulfide bridges decreases success rates as well. PTM sites can indicate accessibility of a residue explaining the higher fraction of modification sites in successful immunogens. However, the actual attachment of a modification might conceal the epitope, especially considering that HPA immunogens were produced in *E.coli*^18^ and thus might lack PTMs present in mammalian proteins leading to differences between immunogen and antibody target.

The HPA dataset was an invaluable resource for this study. The HPA’s extensive coverage of the human proteome, the standardized methodology for antibody production and validation, and the inclusion of negative results provided a robust foundation. However, multiple drawbacks of this dataset have to be contemplated as well.

One notable issue stems from the length of the utilized immunogens of the HPA, termed protein epitope signature tags (PrESTs). With lengths often between 100 and 150 amino acids^18^, these immunogens do not exhibit the properties of peptides and will certainly contain more than one epitope^22^ increasing the complexity of their analysis. While we limited the dataset to immunogens of 50 amino acids or less, it is still highly likely that several epitopes exist in these immunogens. The polyclonal HPA antibodies this analysis is based on will bind different parts of the immunogen in an unknown ratio which can impact performance immensely. This layer of uncertainty might lessen the strength of the differences observed between inadequate and suitable immunogen sequences.

Furthermore, the included antibody-based technologies utilize denaturation and epitope retrieval steps^18^. Those processes lead to the proteins to be detected to change their conformation or unfold^32^. While this step can improve performance of antibodies that were raised against protein fragments, it limits our insights into the antibody’s performance to detect the native protein which is the basis of immunoassays. We hypothesize that many of the trends observed in our analysis would be more pronounced if the antibodies were evaluated regarding their performance in native protein detection. In this case, the correct selection of the immunogen fragment in regard to structural properties would be even more vital^9^ as buried epitopes and differences in secondary structures between immunogen and antibody target would be a larger obstacle. The availability of antibody performance results in technologies detecting native proteins, e.g., ELISA, would significantly enhance this type of study.

Another issue to consider is falsely categorized antibodies and immunogens. Antibodies can be deemed “uncertain” within the HPA if the localization of proteins does not agree with the literature^20^. However, this assignment might also be caused by missing literature annotations regarding the protein’s expected location instead of a wrongly binding antibody. Further, WBs were defined as unspecific if bands appear that indicate an unexpected molecular weight. This definition might again be a false negative as molecular weight can be influenced by multiple factors, e.g., PTMs or dimer formation^33^. An unexpected molecular weight for a protein might thus again be the result of missing knowledge about this protein and its proteoforms. This ambiguity is the reason the HPA classifies these results as “uncertain”.

Our findings underscore the importance of reporting and publishing negative or uncertain data^34^. Negative results are invaluable for understanding the limitations and potential pitfalls in all research, but are especially vital in improving immunogen and antibody selection. By sharing negative data, researchers can contribute to a more comprehensive and realistic understanding of antibody performance, ultimately leading to better guidelines and improved antibody development processes.

In conclusion, our analysis highlights the necessity of systematic investigations into immunogen and antibody choices. Our rigorous approach, supported by the robust data from the HPA, to define successful and unsuccessful antibodies and to evaluate a wide range of structural and functional features confirmed and expanded strongly needed recommendations for researchers working with antibody-based technologies. To the best of our knowledge this is the first study using the available data on HPA antibodies to gain these valuable insights for the scientific community. This study emphasizes the need for data-driven guidelines and the importance of considering both positive and negative data in the pursuit of the generation of more effective and reliable antibodies. Future studies focusing on antibody-based protein detection in native conditions will further enhance our understanding and refine the selection processes for immunogens and antibodies.

## Methods

### Data collection

Information on both successful and unsuccessful antibodies as well as their associated immunogen sequences was required to compare relevant properties. The HPA (www.proteinatlas.org) contains standardized antibody validation results and in the case of in-house produced “HPA antibodies” the utilized immunogen sequence is also available.

Relevant data of the HPA can be accessed via the XML webpages for each entry, i.e., protein. On 10 August 2023, the Ensembl IDs included in version 23.0 of the atlas were used to parse the XML pages and extract the relevant information for each protein and its associated HPA antibodies. For each antibody the outcome of the antibody validation experiments (uncertain, approved, supported or enhanced) in IHC, ICC, and WB were collected. For WB results we also extracted the comments on detected bands if these were available. For 16.510 human proteins with a UniProt ID, 23.616 antibodies with a known immunogen sequence and at least one validation were available. Note that a part of the WB results is not accessible from the XML webpages and could thus not be included in this study.

### Antibody classification

The HPA provides a scale to describe the results of antibody validation experiments of IHC, ICC and WB. For immunogen and antibody analysis, definitions were simplified: an “uncertain” result indicates that the antibody was unsuccessful. An “approved”, “supported”, or “enhanced” result indicates that the antibody was successful for this specific technology. In this study, an antibody and its immunogen are defined as “successful” if the validation results were successful for all three technologies or if the antibody was not tested for one of the technologies but successful in the two remaining technologies. Conversely, an antibody and its immunogen are defined as “unsuccessful” if the validation failed for all three technologies or if the antibody was only tested in two of the technologies and failed for both.

The HPA provides additional information on the results of the WB experiments. Explicitly, the specificity of an antibody was evaluated by including comments on the observed bands. Thus, antibodies with WB results were further classified into specific, unspecific and unsuccessful groups. Specific WBs detected a single band corresponding to the predicted size in kDa (+/- 20%). Unspecific WBs showed a band differing more than +/-20% from predicted size based on the amino acid sequence while the band at the predicted size could either be present or absent. WBs detecting no bands were classified as “no binding”.

### Immunogen analysis

To allow a trade-off between data availability and epitope characterization, only immunogens with a length of 50 amino acids or less were included for the immunogen-specific analysis. To exclude outliers that might skew the analysis, we also filtered for proteins of less than 100 or more than 2000 amino acids in length. These definitions led to 815 successful and 188 unsuccessful antibodies in our final dataset.

After creation and filtering of the data, the final set of immunogens was analysed to infer systematic differences between successful and unsuccessful immunogens that can guide future immunogen and antibody selection. Each immunogen was mapped to the respective full protein sequence to determine its position. HPA immunogens were classified regarding their position within the protein. “Terminus” immunogens include all immunogens which are located fully or partly within the first or last 25 residues of the full-length protein sequence. All other immunogens are considered “center” immunogens.

### Structural immunogen analysis

Predicted protein structures were downloaded from the AlphaFold Protein Structure Database in the format of PDB files^27^. AlphaFold models are currently the state of the art regarding protein structure prediction^35^, thus the predicted models are expected to be highly accurate. The DSSP algorithm was applied^36^ to assign secondary structures (coil, helix or sheet) and surface accessibility to each protein residue. Residues were labeled as buried if their surface accessibility within the protein structure was less than 7% of their potential maximum accessibility. For each immunogen residue, the number of intra-chain contacts were calculated by counting any atom of one residue being within 8 Å of any atom of another residue. These counts were than averaged per immunogen. NetSurfP-2.0 results of disorder prediction were utilized for disorder analysis of immunogens^37^. “High disorder” immunogen were comprised of only residues with a disorder score of at least 0.8 or higher. Vice versa, “low disorder” immunogens contain only residues with a disorder score of 0.2 or lower. All other immunogens were defined as “medium disorder”.

### Functional immunogen analysis

Functional annotations for transmembrane residues, disulfide bridge positions and PTM sites (i.e., annotations of modified residues, glycosylation and lipidation) were included from UniProt^38^.

### Subcellular localization predictions

Annotations of the predicted subcellular localization of all human proteins were available from previous work^39^ and are based on results of DeepLoc^40^.

### R package development

The R package *immunogenViewer* accesses available information on protein structural and functional features from the UniProt database^38^ and the PredictProtein webserver^41^. For extensive information and tutorials of the package, please refer to the documentation (https://bioconductor.org/packages/devel/bioc/html/immunogenViewer.html). In brief, the protein features are visualized along the sequence of the protein of interest and immunogens can be added and visualized as well.

### Statistical testing

Testing for statistical significance was performed using the Mann-Whitney U test or Chi-squared test. The cutoff for significance was set at .05.

## Author contributions

KW conceptualized the study, was involved in data curation, formal analysis, investigation, methodology, software and visualization, and wrote the original draft. HK was involved in conceptualization, supervision, and review and editing of the manuscript. KB and HZ were involved in review and editing of the manuscript. CET was involved in funding acquisition and review and editing of the manuscript. SA was involved in conceptualization, supervision and review and editing of the manuscript. All authors read and approved the final manuscript.

## Acknowledgements

We would like to acknowledge the support received from the Netherlands eScience Center during software development. Especially the valuable feedback and knowledge shared by research software engineers Giulia Crocioni and Dr. Pablo Rodríguez-Sánchez was appreciated as well as the financial support KW received through the eScience fellowship program 2023/2024. KW was kindly supported by Alzheimer Nederland. KW, HZ, CET, and SA are supported by the European Union’s Horizon 2020 research and innovation programme under the Marie Skłodowska-Curie grant agreement No 860197 (MIRIADE). KB is supported by the Swedish Research Council (#2017-00915 and #2022-00732), the Swedish Alzheimer Foundation (#AF-930351, #AF-939721, #AF-968270, and #AF-994551), Hjärnfonden, Sweden (#FO2017-0243 and #ALZ2022-0006), the Swedish state under the agreement between the Swedish government and the County Councils, the ALF-agreement (#ALFGBG-715986 and #ALFGBG-965240), the European Union Joint Program for Neurodegenerative Disorders (JPND2019-466-236), the Alzheimer’s Association 2021 Zenith Award (ZEN-21-848495), the Alzheimer’s Association 2022-2025 Grant (SG-23-1038904 QC), La Fondation Recherche Alzheimer (FRA), Paris, France, the Kirsten and Freddy Johansen Foundation, Copenhagen, Denmark, and Familjen Rönströms Stiftelse, Stockholm, Sweden. HZ is a Wallenberg Scholar and a Distinguished Professor at the Swedish Research Council supported by grants from the Swedish Research Council (#2023-00356; #2022-01018 and #2019-02397), the European Union’s Horizon Europe research and innovation programme under grant agreement No 101053962, Swedish State Support for Clinical Research (#ALFGBG-71320), the Alzheimer Drug Discovery Foundation (ADDF), USA (#201809-2016862), the AD Strategic Fund and the Alzheimer’s Association (#ADSF-21-831376-C, #ADSF-21-831381-C, #ADSF-21-831377-C, and #ADSF-24-1284328-C), the European Partnership on Metrology, co-financed from the European Union’s Horizon Europe Research and Innovation Programme and by the Participating States (NEuroBioStand, #22HLT07), the Bluefield Project, Cure Alzheimer’s Fund, the Olav Thon Foundation, the Erling-Persson Family Foundation, Familjen Rönströms Stiftelse, Stiftelsen för Gamla Tjänarinnor, Hjärnfonden, Sweden (#FO2022-0270), the European Union Joint Programme – Neurodegenerative Disease Research (JPND2021-00694), the National Institute for Health and Care Research University College London Hospitals Biomedical Research Centre, and the UK Dementia Research Institute at UCL (UKDRI-1003). Research of CET is supported by Innovative Medicines Initiatives 3 TR (Horizon 2020, grant no 831434) EPND (IMI 2 Joint Undertaking (JU), grant No. 101034344) and JPND (bPRIDE), National MS Society (Progressive MS alliance), Alzheimer’s Drug Discovery Foundation, Alzheimer’s Association, Health Holland, the Dutch Research Council (ZonMW), Alzheimer’s Drug Discovery Foundation, The Selfridges Group Foundation, Alzheimer Nederland. CT is recipient of ABOARD, which is a public-private partnership receiving funding from ZonMw (#73305095007) and Health∼Holland, Top Sector Life Sciences & Health (PPP-allowance; #LSHM20106). CET is recipient of TAP-dementia, a ZonMw funded project (#10510032120003) in the context of the Dutch National Dementia Strategy.

