## Supplementary figures for "Data-driven evaluation of suitable immunogens for improved antibody selection"

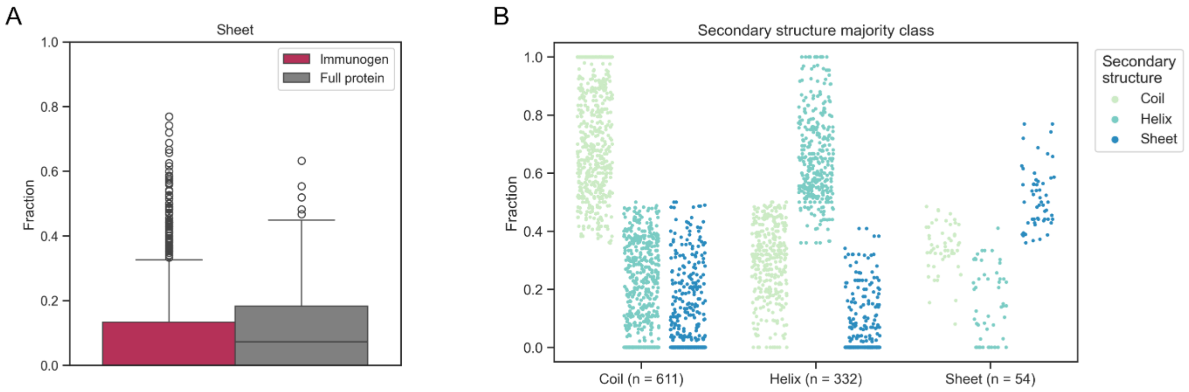


**Figure S1. Sheet content in HPA immunogens.** Compared to the full protein sequences of the HPA dataset, the selected immunogen regions are depleted of sheet residues (A). Most immunogens are dominated by coil residues. The small set of immunogens in which sheets are the majority secondary structure still contains fractions of coils and helices (B). HPA - Human Protein Atlas
